# Linked selection of insertions and deletions in coding regions of the great tit genome

**DOI:** 10.1101/2025.09.26.678803

**Authors:** Yu-Chi Chen, Justin J.S. Wilcox, Toni I. Gossmann

## Abstract

Insertions and deletions (indels) are a ubiquitous form of genomic variation; yet, their contribution to adaptive evolution remains poorly understood. Technical challenges in identifying indels and integrating them into existing evolutionary models have limited their study, but previous work suggests that they may be subject to strong selective pressures. Here, we investigated selective signatures of indels in coding regions of the great tit (*Parus major*). We identified species-specific indels in protein-coding genes and analyzed the 300 bp flanking regions of indels using codeml to estimate dN, dS, and *ω* (dN/dS). Additionally, we performed enrichment analyses and protein structure modeling to assess functional consequences. We found that mutations in the flanking regions of indels are non-random and that selection strength varies with indel type and length. Genes including indels were enriched for functions related to protein and nucleic acid binding and linked to potentially adaptive traits such as vocal learning, migration, circadian rhythm, and environmental acclimation. Our results suggest that insertions and deletions may serve as hotspots of selection, influencing nearby sequences and contributing to adaptation in the great tit.

**Significance Statement:** Small DNA changes called insertions and deletions (indels) are common in genomes, but their role in evolution is often overlooked because they are difficult to study. Using the genome of the great tit, a well-studied songbird, we identified thousands of indels in protein-coding regions and examined how they affect nearby DNA. We found that indels are not neutral accidents: they show signs of natural selection and may influence important traits. Some of the affected genes are linked to behaviors and adaptations that are crucial for survival, such as vocal learning, migration, and adaptation to new environments. These results suggest that indels can act as hotspots for evolutionary change and may help explain how species adapt to their surroundings. By highlighting the evolutionary importance of these often-neglected mutations, our study opens new perspectives on the genetic basis of adaptation.

## Introduction

Mutations and selection shape the evolutionary process. Mutations introduce genetic variations and fuel genomic diversity, while selection acts on these variations to shape adaptive traits. With the advancement of bioinformatic and sequencing technologies, genetic variation has become much easier to identify. Such variation is widely used in evolutionary, population, and functional genomics. However, the influences of selection on genetic variation are still incompletely understood, leaving gaps in our understanding of the major forces shaping evolution. Specifically, most studies have focused on single nucleotide polymorphisms (SNPs), often overlooking other forms of variations, such as insertions and deletions (indels).

Indels, which have been studied far less extensively than SNPs in evolutionary and functional genomics, can range in length from a single base to several thousand bases [1, 2]. Small indels are typically referred to simply as indels, whereas large indels are classified into structural variations (SVs). A commonly used cutoff between small and large indels is 50 bp [2, 3, 4, 5]; however, it is worth noting that this threshold is arbitrary and may range from 4 to 10,000 bp in different studies [6, 7, 8, 9, 1, 10, 11].

Despite this arbitrary cutoff, small and large indels may arise through different molecular mechanisms and exhibit distinct characteristics. Small indels are often generated during DNA double-strand break (DSB) repair processes such as non-homologous end joining (NHEJ), microhomology-mediated end joining (MMEJ), and single-strand annealing (SSA) [12, 13]. While large indels can arise from DSB repair [13], they are also associated with transposon activity and other large-scale genomic rearrangements [14, 15]. As SVs, large indels are notable for their highly heterogeneous mutation rate [16]. Although large indels are much less abundant than small indels [10, 17, 7, 18], they have greater potential to change protein structures and functions due to their greater sequence length. These mechanistic and feature differences suggest that small and large indels may be subject to different evolutionary dynamics and selective pressures. These differences suggest that small and large indels may be subject to distinct evolutionary dynamics and selective pressures.

While these features make indel detection technically challenging, for example during variant calling [19, 20, 21], indels can alter the structure and length of DNA and proteins. Such changes, which SNPs cannot capture, suggest that indels may have substantial fitness effects on DNA and protein function and can uniquely influence evolutionary trajectories. Consequently, indels may contribute to adaptive and phenotypic variation that cannot be explained by SNPs alone. At the molecular level, indels can directly change DNA dosage and genomic structure, thereby altering gene expression and/or impairing gene functions [22, 23, 24]. This direct effect implies that indels are often deleterious and constrain adaptation, but also makes indels more likely to affect phenotypes than SNPs [25, 26, 27, 28, 29]. At the population level, indels influence divergence and speciation. They can influence fitness by suppressing recombination and reducing the efficacy of purifying selection. [29, 30, 31, 32]. Together, these direct and indirect effects suggest that indels may play pivotal roles in functional changes of DNA/protein sequences and in population diversification.

Although indels can influence molecular function and population diversity, their contribution to sequence divergence and interaction with selection remains unclear. In modern humans, indels contribute more to sequence divergence than SNPs [1, 33], and they can better clarify past evolutionary events because they are often under stronger selective forces [26, 34]. However, human specific-indels contribute less to sequence divergence in the coding regions than in introns, both in number and in length[35]. This suggests that most coding-region indels are subject to strong purifying selection. Despite this constraint, indels have been implicated in adaptive traits [36, 37, 38], indicating that a subset may be positively selected.

The selective forces acting on indels depend on multiple factors. Although insertions and deletions are often grouped together, they represent two distinct types of mutations with different molecular characteristics and evolutionary patterns. Indels are more common in GC-rich regions; however, only regions near insertions are enriched with CpGs [39, 25]. In coding regions, deletions are subject to stronger purifying selection than insertions [40, 26, 27]. Even so, deletions are more abundant than insertions [26]. Other than indel type, indel length can also affect gene expression and thereby influence selection strength in the coding regions [24]. Taken together, indel type and length both modulate the selective forces acting on them.

As indels are under strong selective forces, they may influence their flanking regions through linked selection. Previous research reported increased nucleotide divergence between closely related species in flanking regions of indels, which gradually decreases with distance over several hundred bases. This pattern coincides with decreased transition substitutions and increased transversion substitutions in flanking regions of indels compared to regions further away from them [41, 42], suggesting selective pressures in indel-flanking regions. Furthermore, indels occur more frequently in GC-rich regions [39]. More broadly, CpG distribution has been shown to shape mutational dynamics and selective pressures in protein-coding DNA across vertebrates [43]. These observations imply that flanking regions of indels may act as hotspots of selection.

The great tit genome is an excellent model for studying selective pressures [44, 45, 46, 47]. Avian genomes are relatively small, syntenic, and computationally tractable, and the great tit reference genome is high-coverage and well annotated [45, 48]. Moreover, previous work showed that indels in the great tit are under selection [27]. Estimates of the distribution of fitness effects (DFE) suggest that great tit-specific indels experience stronger purifying selection in protein-coding regions than SNPs. In addition, noncoding indels are associated with reduced nucleotide diversity, hinting at linked selection [27]. In contrast, linked selection around coding-region indels remains poorly understood. The depth of genomic resources available for the great tit makes it well suited for examining this phenomenon.

Indels may play a pivotal role in species evolution and adaptation. They can be under selection and may have substantially different impacts on flanking regions, yet direct evidence of these selective pressures is limited. In our study, we use the great tit as a model species to investigate how selection acts on indels and their flanking regions. Specifically, we (i) identified great tit-specific indels in coding genes, (ii) examined selective signatures (dS, dN, and *ω*) in the flanking regions, (iii) compared selection across different indel types (insertions vs. deletions) and lengths (small *<* 50 bp vs. large ≥ 50 bp), and (iv) tested whether genes harboring indels are enriched for biological functions relevant to adaptation. This framework allows us to assess the evolutionary impact of indels on protein-coding regions in the great tit.

## Materials and Methods

### The dataset

We obtained the genome assemblies for the great tit (*Parus major*) (RefSeq ID: GCF_001522545.3) and two closely related avian species with high-quality reference genomes, the collared flycatcher (*Ficedula albicollis*) (RefSeq ID: GCF_000247815.1) and the zebra finch (*Taeniopygia guttata*) (RefSeq ID: GCF_003957565.2) from the NCBI RefSeq database [49] via ftp. The hooded crow (*Corvus cornix*) (RefSeq ID: GCF_000738735.6) was used as the outgroup in all analyses. All data were retrieved before August 16, 2024.

### Multispecies alignment

Our pipeline is shown in Figure 1 A. For each protein-coding gene, we extracted the longest isoform and obtained the corresponding nucleotide and amino acid sequences for each focal species using AGAT v0.8.0 [50] (Figure 1 A, step 1). The nucleotide sequences were clustered into orthogroups with OrthoFinder v2.5.5 using the blast_nucl option [51] and single-copy orthogroups were retained for alignment (Figure 1 A, step 2). Amino acid sequences within each orthogroup were aligned with PRANK v.170427 [52] with the “permanent insertions” option (+F) and a guide tree based on the avian phylogeny reported by Shaohong Feng et al [53] (Figure 1 A, step 3).

**Figure 1:**
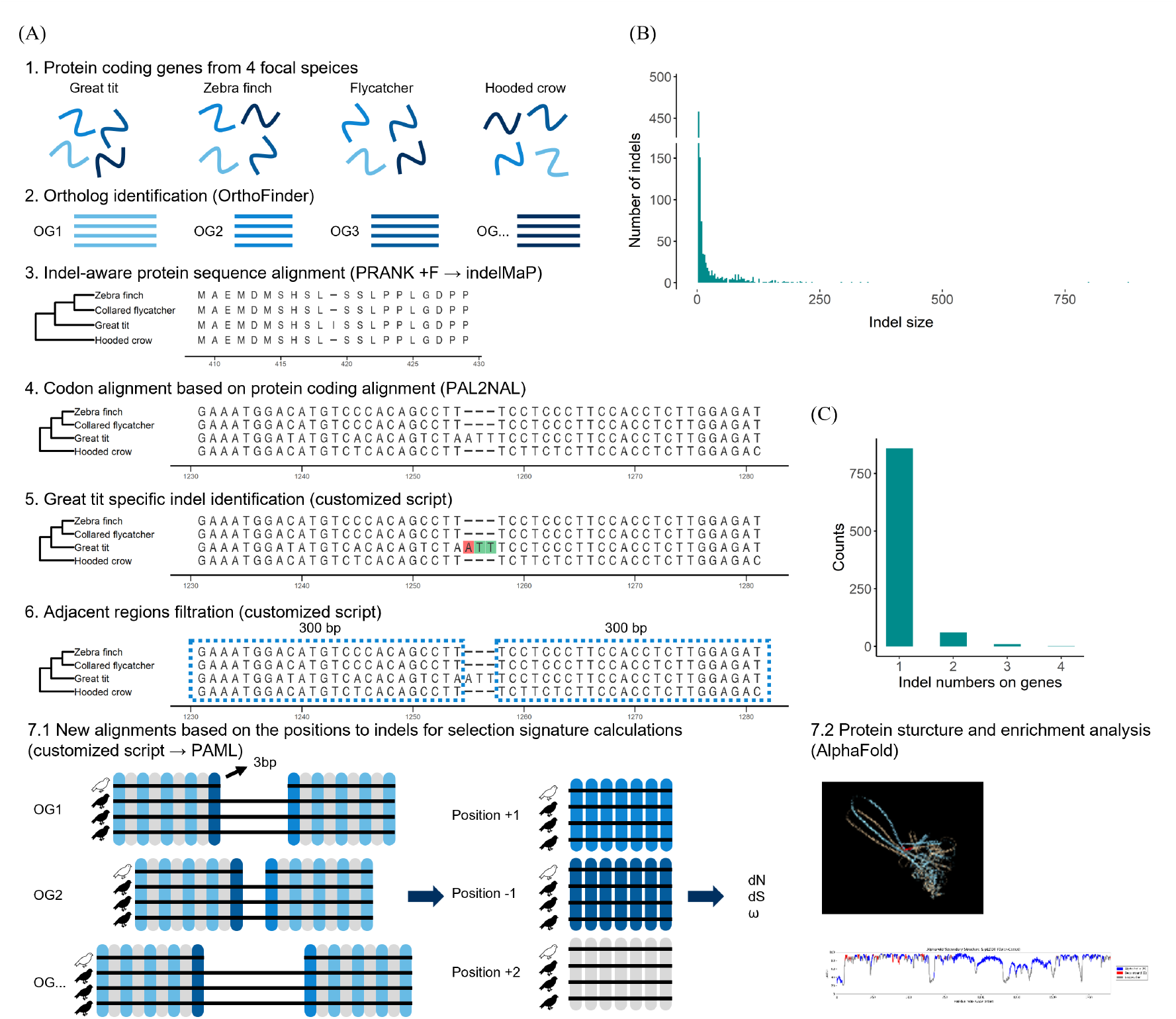
Workflow and summary statistics for indel identification. **(A)** Pipeline for examining selection signatures in regions adjacent to great tit–specific indels. **(B)** Length distribution of great tit–specific indels identified by the pipeline. **(C)** Number of great tit–specific indels per orthogroup alignment.

### Indel identification

Before identifying the great tit-specific indels, we first reconstructed ancestral sequences in each alignment using indelMaP [54] (Figure 1 A, step 3). Alignments were excluded if any sequence was out of frame after this step. The remaining amino acid alignments were converted into codon alignments with PAL2NAL v14 [55] using default settings (Figure 1 A, step 4). Custom scripts were then applied to filter gaps according to the following criteria for great tit-specific indel events: (1) gaps present in the lowest common ancestor of the great tit and closely related species but absent in the great tit were classified as great tit-specific insertions; (2) gaps present in the great tit but absent in the lowest common ancestor of the great tit and closely related species were classified as great tit-specific deletions. In other words, great tit–specific insertions are regions uniquely present in the great tit, whereas great tit–specific deletions are regions uniquely absent in the great tit (Figure 1 A, step 5).

### Selection tests

To evaluate selection around indels, we defined the upstream and downstream flanking sequences (300 bp each) as flanking regions of indels. Indels were excluded if their flanking regions overlapped with gaps from other species within this 300 bp window (Figure 1 A, step 6). To prevent alignment errors at sequence termini, flanking regions shorter than 3 bp were also removed. Great tit-specific indels were grouped into three sets: (i) all indels combined, (ii) small indels (*<* 50 bp), and (iii) large indels (≥ 50 bp). Flanking sequences were extracted for selection analysis using bedtools v2.31.1 [56].

Selection signatures were estimated by concatenating codons according to their position relative to indels and aligning them across species. We calculated the non-synonymous substitution rate (dN), the synonymous substitution rate (dS), and the nonsynonymous/synonymous substitution ratio (*ω*) with CODEML in PAML [57]. Three branch models were applied: (1) free-ratios model (model = 1, NSsites = 0, omega = 2, fix_omega = 0), (2) two-ratio model with the great tit as the foreground branch (model = 2, NSsites = 0, omega = 2, fix_omega = 0) (Figure 1 A, step 7.1).

### Enrichment analysis

For each indel set, we extracted the corresponding genes and performed Gene Ontology (GO) annotation and enrichment analysis using DAVID [58, 59] and protein-protein interaction analysis using STRING [60]. To minimize bias, the longest isoforms of each protein-coding gene was used as the background set (Figure 1 A, step 7.2).

### 3D protein structure analysis

For selected genes harboring great tit-specific indels with previously reported functions in birds, we predicted protein structures using the AlphaFold Server [61]. The structural impact of the indel-containing regions was assessed by comparison with orthologous proteins from the hooded crow. Predicted structures were visualized with UCSF ChimeraX [62] (Figure 1 A, step 7.2).

## Results

### Indel identification

We searched for indels in protein-coding regions of our four focal species, the great tit, collared flycatcher, zebra finch, and hooded crow, using alignments of the longest isoforms from each species. We identified 15,120 aligned orthogroups of protein coding genes in these species, of which 12,421 were single-copy orthogroups. After aligning the genes from these single-copy orthogroups, only 27 alignments (0.22% of all the single-copy orthogroups) contained sequences that were not in frame. These frameshift alignments were excluded. We further identified 2,134 orthogroups without gaps between species, which served as the controls in the selection test. Approximately 80% of the orthogroups contained gaps, but most were excluded because the indels or their flanking regions overlapped with other indels or other flanking regions. After filtering, we retained 1,015 high-confidence great tit-specific indels located within 931 orthogroups. Among these orthogroups, 92.27% contained a single great tit-specific indel, and only one orthogroup contained 4 indels (Figure 1 C). Of the 1,015 indels, 634 (62.50%) were deletions and 37.50% were insertions (Table 1). The indel lengths ranged from three to 879 bp, with a mean of 23.65 bp. The vast majority (93.98%) were small indels (*<* 50 bp; 875 indels), while only 140 were large indels (≥ 50 bp) (Figure 1 B).

**Table 1:** Counts of great tit–specific indels with non-overlapping 300 bp flanks.

|  |  |
| --- | --- |
| Flanking region length | 300 bp |
| Great tit–specific indels (total) | 1,015 |
| Insertions | 381 |
| Deletions | 634 |
| Indel length ≥ 50 bp | 140 |
| Indel length $<$ 50 bp | 875 |

### Selection signatures in flanking regions of indels

We evaluated indels using three substitution signatures (non-synonymous substitution rate, dN; synonymous substitution rate, dS; *ω*) revealed distinct patterns across the flanking regions of indels under the two ratio model. The mean dN of indels was 0.014 *±* 0.0061, significantly higher than the control value calculated from the 2,134 gap-free alignments (mean_control_ = 0.0044 *±* 0.0010; *t* -test, *p <* 2.2e-16). The values peaked three codons away from the indel sites (site 0), reaching nearly four times higher than the values at more distant sites. This elevated dN near indels persisted regardless of subgrouping (Figure 2).

**Figure 2:**
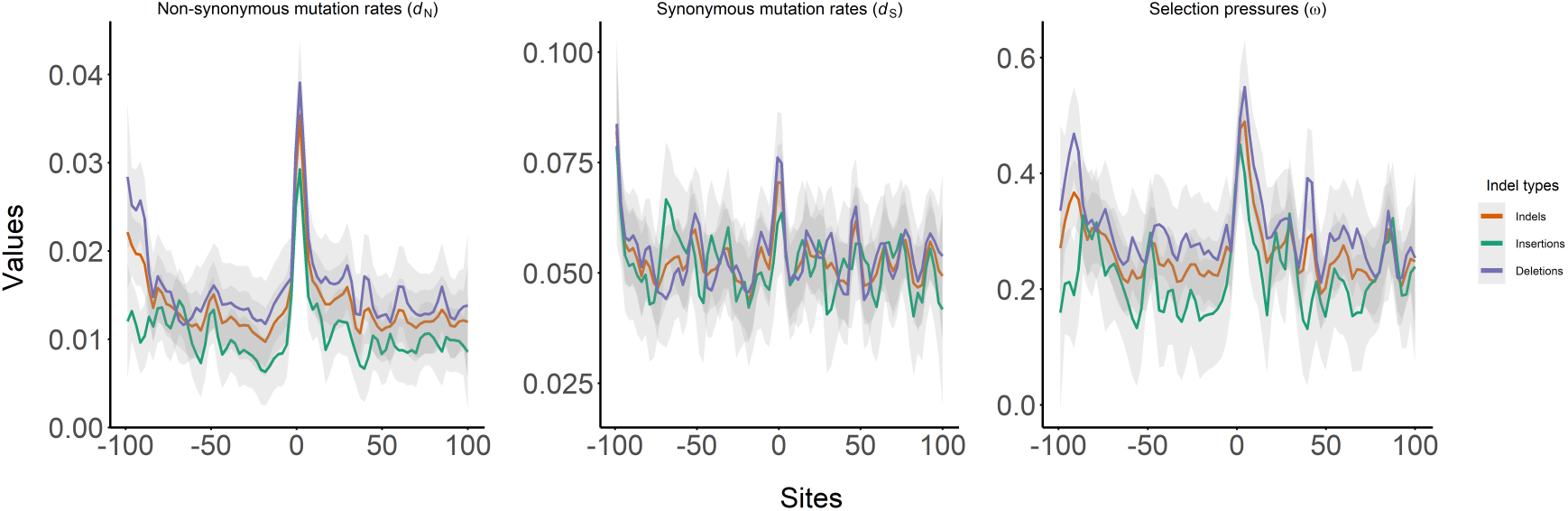
Spline plots of selection signatures, dN, dS, and *ω* within 300 bp flanking regions of the great tit-specific indels (orange), great tit-specific insertions (green), and great tit specific deletions (purple) under the two-ratio branch model. Curves show the non-synonymous rate (dN), synonymous rate (dS), and dN/dS (*ω*) by codon position relative to the indel (site 0).

The mean dS was 0.054 *±* 0.022 [SD], also significantly higher than the control (mean_control_ = 0.048 *±* 0.0062; *t* -test, *p* = 0.000306); however, unlike dN, dS values were relatively stable across distances from indels. The mean *ω* for indels was 0.27 *±* 0.084, significantly higher than the control (mean_control_ = 0.092 *±* 0.023; *t* -test, *p <* 2.2e-16). Peaks in *ω* occurred immediately adjacent to indels, reaching about twice the values observed at more distant sites (Figure 2).

### Effect of indel length on selection signatures

When subgrouping indels by lengths, both small (*<* 50 bp) and large (≥ 50 bp) indels showed dN peaks close to the indel site. For small indels, the mean dS (0.054 *±* 0.021) did not significantly differ from all indels combined (*t* -test, *p* = 0.9543). However, the mean dN (0.015 *±* 0.0065) and the *ω* (0.29 *±* 0.097) were significantly higher than in the pooled dataset (*t* -tests, *p* = 0.0440 and 0.00743, respectively). For large indels, the mean dS (0.054 *±* 0.043) was similar to that of all indels (*t* -test, *p* = 0.9608). However, the mean dN (0.0063 *±* 0.0066) were significantly lower, and the mean *ω* (20.14 *±* 139.83) were significantly higher but with large SD (*t* -tests, *p <* 2.2e-16 and 0.04636, respectively) (Figure 3).

**Figure 3:**
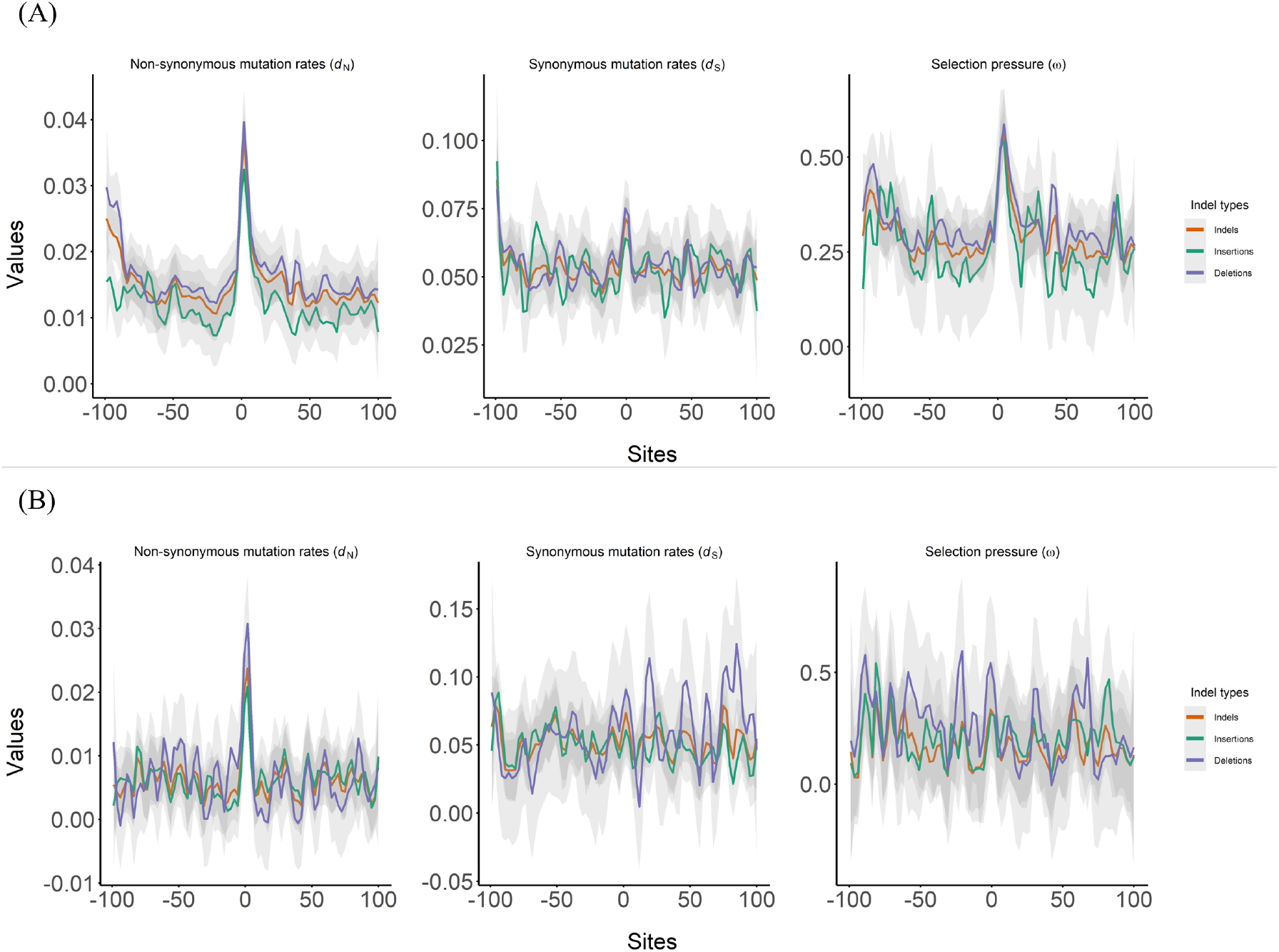
Selection signatures within 300 bp flanks by indel length category. **(A)** Small indels (*<* 50 bp). **(B)** Large indels (≥ 50 bp). Values greater than 1.00 for *ω* are capped at 1.00 for display.

When comparing indel length categories, small indels had significantly higher dN (0.015 *±* 0.0065) than large indels (0.0063 *±* 0.0066; *t* -test, *p <* 2.2e-16). Mean dS did not differ significantly (0.054 *±* 0.021 vs. 0.054 *±* 0.043; *t* -test, *p* = 0.9313). Mean *ω* was significantly higher in small indels (0.29 *±* 0.097) than in large indels (20.14 *±* 139.83; *t* -test, *p* = 0.04663) (Figure 3).

### Differences between insertions and deletions

Indel type also influenced selection signatures. We categorized great tit–specific indels into two polarities: great tit–specific insertions, where an inserted region is unique to the great tit, and great tit–specific deletions, where deleted regions are unique to the great tit. This is effectively polarization by parsimony. When we compared the selection signatures, the mean dN for insertions (0.011 *±* 0.0055) was significantly lower than for deletions (0.016 *±* 0.0074; *t* -test, *p* = 3.716e-14). By contrast, mean dS did not differ significantly between insertions (0.053 *±* 0.027) and deletions (0.055 *±* 0.023; *t* -test, *p* = 0.4693). Mean *ω* followed the same pattern as dN, with higher values for deletions (0.30 *±* 0.11) than insertions (0.22 *±* 0.12; *t* -test, *p* = 1.438e-11) (Figure 3).

Codon and amino acid frequency analyses revealed different selective preferences between insertions and deletions. We calculated the ratio of codon and amino acid counts in the inserted and deleted regions to their respective counts in the longest isoform sequences of all protein-coding genes in the great tit. The most frequently inserted codons were TGT, TCT, and ACT, while the least frequent were CGC, ATC, and CCG. In contrast, deleted codons were more evenly distributed across all codons (Figure 4B). At the amino acid level, serine (S) was the most deleted residue and leucine (L) was the most inserted residue, while methionine (M) was the least frequent in both categories (Figure 4A).

**Figure 4:**
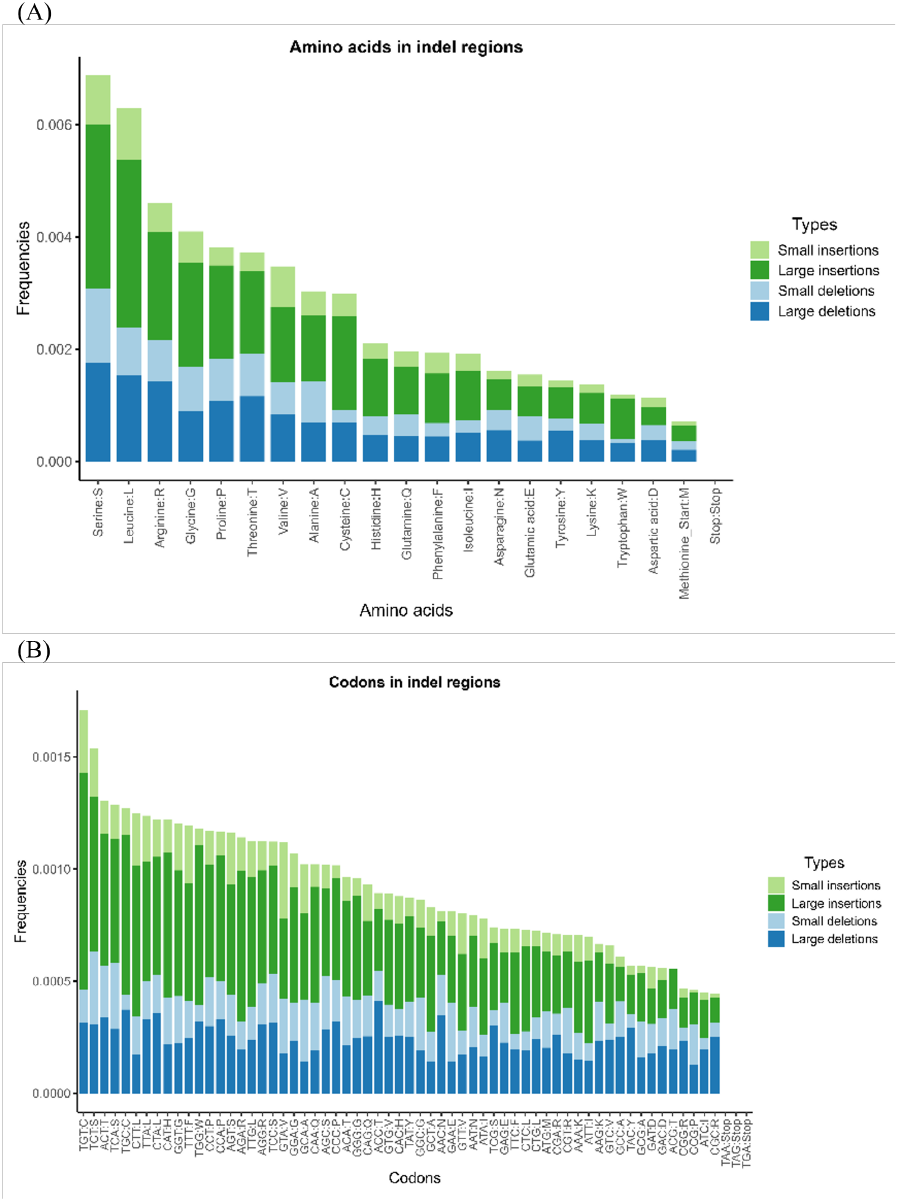
Frequencies of inserted and deleted sequence content in great tit–specific indels. **(A)** Amino acids. **(B)** Codons.

### Functional analysis of indels

Genes harboring great tit-specific indels were associated with diverse molecular functions and protein–protein interactions. Gene Ontology (GO) enrichment showed strong overrepresentation of structural and binding functions, including protein binding, DNA binding, RNA binding, and actin filament organization (Figure 5A). Genes with insertions were enriched for metal ion binding, nucleic acid binding, and regulation of transcription by RNA polymerase II, whereas genes with deletions were enriched for protein binding, cell adhesion molecule binding, and DNA-directed DNA polymerase activity. Notably, only deletion-harboring genes showed enrichment in 13 annotation clusters, including the anemia pathway and autophagy (see Supplementary Table 13). In protein–protein interaction networks, only deletion-harboring genes were enriched (e.g., intracellular non-membrane-bounded organelles and coiled-coil proteins).

**Figure 5:**
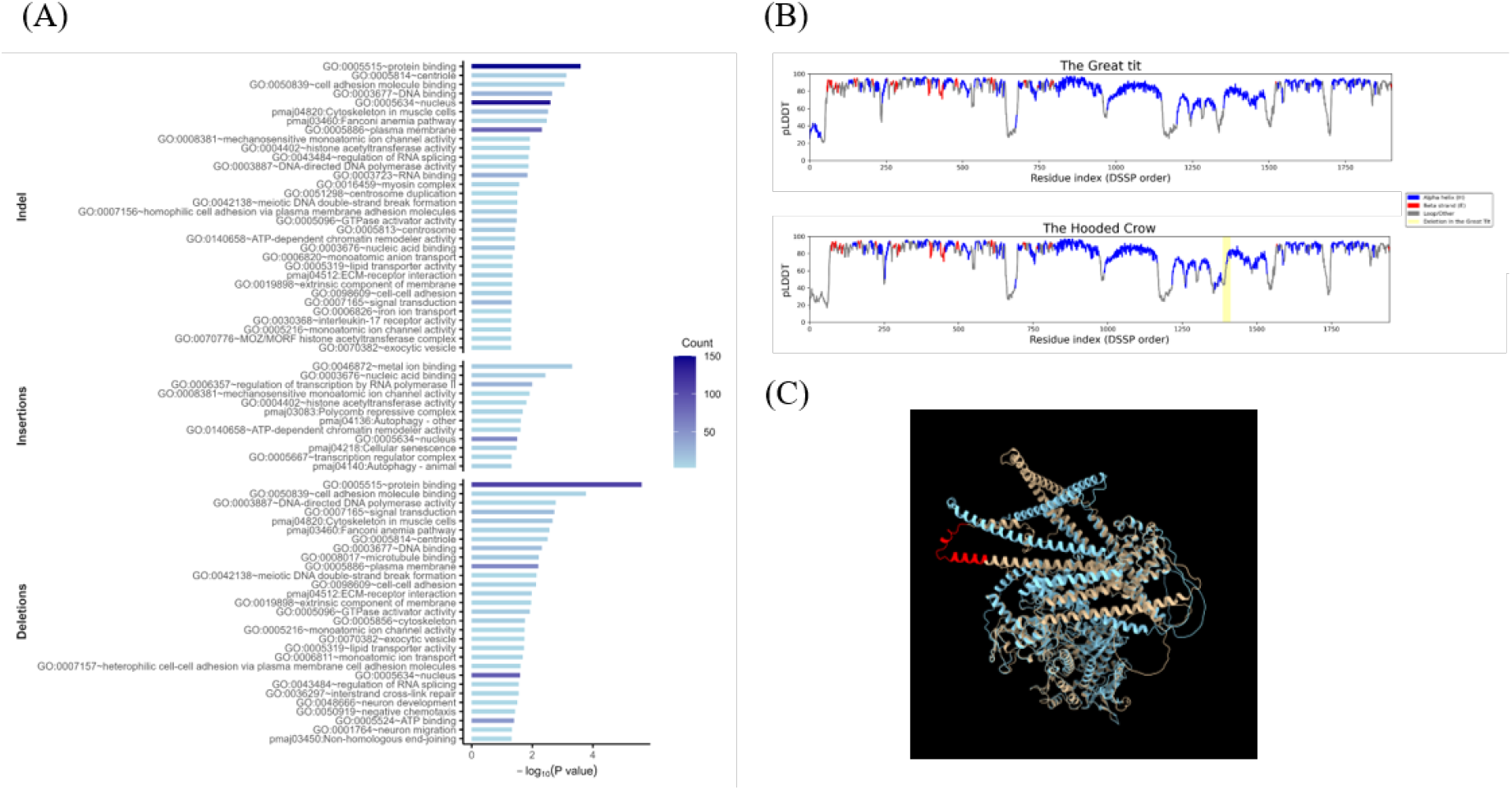
Functional analyses of genes harboring great tit–specific indels. **(A)** GO enrichment for genes with indels. **(B)** Predicted local confidence (pLDDT) for *MYO5A* in great tit and hooded crow. **(C)** AlphaFold-predicted protein structures: great tit (light blue) and hooded crow (light orange); the deleted fragment in great tit is highlighted in red.

Several genes with indels were linked to adaptive traits in the great tit and other birds. These included genes associated with vocal learning (e.g., *ATRX, CACNA1B, CACNA2D3, GRIK2, GRIN2B, LRRC8C, LRRC8D, MMP2, TPCN1, UTS2B*; Table 2), environmental adaptation (*CREBBP, DRD3, DRD4, EP300, GDF9, KAT6A, MAPK13, SPINT1, VAV3*), and migration (*ATC2B, CLOCK, DRD4, SDC1, PER3*). In particular, *CLOCK* and *DRD4* are involved in multiple adaptive traits related to circadian rhythm, including migration and breeding timing.

**Table 2:** Genes harboring great tit–specific indels and associated adaptive traits reported in birds (see cited references).

| Gene | OG_ID | Indel type | Length (bp) | Reported adaptive trait(s) |
| --- | --- | --- | --- | --- |
| ANKRD10 | 7224 | del | 18 | Muscle growth regulation [68] |
| ATG2B | 8941 | del | 3 | Migration [69] |
| ATG2B | 8941 | ins | 3 | Migration [69] |
| ATRX | 7981 | del | 3 | Vocal learning [70] |
| ATRX | 7981 | ins | 3 | Vocal learning [70] |
| CACNA1B | 9771 | del | 60 | Vocal learning [71] |
| CACNA1B | 9771 | del | 3 | Vocal learning [71] |
| CACNA2D3 | 7574 | del | 3 | Vocal learning [71] |
| CD36 | 3549 | del | 3 | Lipid metabolism [72] |
| CLNS1A | 6776 | ins | 3 | Vocal learning [71] |
| CLOCK | 4933 | ins | 3 | Breeding time [73]; circadian clock [74]; adaptation to urbanized areas (methylation) [75]; migration [76, 77] |
| COL4A5 | 7890 | del | 9 | Adaptation to urbanized areas (methylation) [75] |
| COL4A5 | 7890 | ins | 3 | Adaptation to urbanized areas (methylation) [75] |
| CREBBP | 5628 | ins | 30 | Stress response [78]; hypoxia; wide-spread [79] |
| DEAF1 | 9568 | ins | 75 | Epigenetic inheritance [80] |
| DRD3 | 6626 | del | 3 | Adaptation to urbanized areas (methylation) [75] |
| DRD4 | 9567 | ins | 3 | Epigenetic inheritance [80]; circadian clock [81, 74]; adaptation to urbanized areas (methylation) [75]; migration [76, 77] |
| DYSF | 5175 | del | 12 | Muscle growth regulation [82] |
| EP300 | 3903 | ins | 30 | Stress response [78] |
| GDF9 | 12748 | del | 6 | Breeding time [73] |
| GRIK2 | 2099 | del | 3 | Vocal learning [70] |
| GRIN2B | 3864 | del | 9 | Vocal learning [70] |
| IFIH1 | 8249 | del | 6 | Muscle growth regulation [68] |
| KAT6A | 5192 | del | 3 | Hypoxia, wide-spread [79] |
| KAT6A | 5192 | ins | 84 | Hypoxia, wide-spread [79] |
| LRRC8C | 1082 | ins | 3 | Vocal learning [71] |
| LRRC8D | 1083 | del | 3 | Vocal learning [71] |
| MAPK13 | 11096 | ins | 60 | Stress response [78] |
| MMP2 | 10884 | del | 6 | Vocal learning [70] |
| MYH15 | 6560 | ins | 3 | Muscle growth regulation [82] |
| MYO5A | 11216 | del | 81 | Feather pigmentation [63] |
| PER3 | 6162 | ins | 60 | Circadian clock [74, 81]; migration [76] |
| RNF25 | 8203 | del | 9 | Muscle growth regulation [68] |
| SDC1 | 2303 | del | 6 | Migration [69] |
| SPINT1 | 9214 | del | 3 | Hypoxia, wide-spread [79] |
| TPCN1 | 10634 | del | 9 | Vocal learning [71] |
| TRPM3 | 2849 | del | 126 | Ion channels and structural proteins [83, 84] |
| TRPM5 | 9329 | del | 3 | Ion channels and structural proteins [83, 84] |
| TRPM6 | 2857 | ins | 66 | Ion channels and structural proteins [83, 84] |
| UTS2B | 5884 | del | 30 | Vocal learning [85] |
| VAV3 | 1146 | del | 84 | Hypoxia; wide-spread [79] |
| FRMPD4* (FRMPD3) | 7959 | ins | 3 | Vocal learning [70] |

Finally, we examined *MYO5A*, previously implicated in feather pigmentation regulation [63]. The great tit ortholog contained an 81-bp deletion (Figure 5C). Compared with the hooded crow ortholog, this deletion spanned a region linking to the start of an *α*-helix, but did not cause major structural changes in the helix (Figure 5B).

## Discussion

Although indels are the second most common contributors to sequence variation [7], their roles in evolutionary processes and the ways in which selection acts upon them remain insufficiently understood, largely because of technical challenges in detection and analysis. In this study, we identified great tit-specific indels within coding regions and evaluated selection strength indices (dN, dS, and *ω*) in their flanking regions. We applied both the two-ratio and free-ratio models in PAML, and both models produced consistent patterns (see Supplementary Table 1). We further conducted functional analyses of genes harboring great tit-specific indels and examined structural predictions for *MYO5A*, a gene containing a great tit-specific deletion.

Indels are common in coding regions. Our study revealed that even within coding regions, 80% of the alignments contain indels, with the majority of them being in-frame. These results indicate that indels contribute substantially to sequence variation (Table 1) without necessarily exerting deleterious effects, consistent with previous findings in humans [10]. Our filtration criteria are stringent to retain only high-confidence indels for further analysis. Applying our pipeline with shorter flanking regions would likely identify additional species–specific indels for other studies.

Our analyses revealed that indel-flanking regions are subject to stronger selective pressures than control regions, which are located on genes without indels, indicating that selection is elevated near indels compared to non-indel coding regions. Furthermore, even within indel coding regions, selective pressures drop drastically just three bases away from indels (Figure 2). Previous studies have reported that regions adjacent to indels exhibit decreased nucleotide diversity and transversion substitutions, along with increased transition substitutions, relative to regions farther from indels [64, 41], reflecting similar heightened selective pressures observed near indels in our analyses.

Indel length is an important factor influencing selective strength. Previous research primarily focused on whether indel length caused frameshifts and thus affected selective pressures [24], while in our research, our study provides a broader perspective on how indel length impacts selective strength. In great tit, small indels are six times more abundant than large indels (Table 1), which corresponds to previous research in other species [10, 17, 7, 18]. Furthermore, our study demonstrates that selective pressures differ between small and large indels, using the 50 bp cutoff (Figure 3). Small indels are under much stronger positive selection than large indels. Although the cutoff between small and large indels is arbitrary, their distinct selective signature patterns support the idea that they are subject to different evolutionary dynamics and selective pressures.

Our results suggest that the selective forces act differently on insertions and deletions (Figure 2). Previous research showed that deletions are under stronger purifying selection but still more abundant than insertions [40, 26, 27]. Our findings revealed that deletions arise more frequently than insertions but might be largely purged as deleterious [65], with only beneficial deletions retained in functional regions since they are stronger under positive selection. An alternative explanation is insertion-biased gene conversion, whereby high recombination regions exhibit an increased fixation rate of insertions but a reduced fixation rate of deletions [66, 18, 65], which produce patterns that resemble positive selection [67].

The codons that are inserted and deleted occurred at different frequencies. Deleted codons are evenly distributed across all codons, indicating no preference for which codons are removed, whereas certain codons, such as TGT, are inserted more frequently than others (Figure 4B). These patterns suggest distinct selective pressures acting on insertions versus deletions. In contrast, the amino acids that are inserted and deleted reflect general patterns of abundance within the great tit (Figure 4B). Amino-acid insertion and removal frequencies are, however, confounded by codon degeneracy, for which we do not control. For example, serine is encoded by 6 different codons, whereas methionine is encoded by 1 codon. As such, amino acids do not differ in insertion and deletion frequency, but even though codons do. This may reflect some broader interplay between the standard eukaryotic genetic code, codon-usage, and mutational patterns.

The genes harbouring great tit-specific indels are implicated in adaptive traits. The great tit is a widespread passerine, which is currently undergoing pressure from urbanization. We found genes from previous research that support the traits related to migration, vocal learning, stress response, and adaptation to urbanization (Table 2). Interestingly, we found two genes, *CLOCK* and *DRD4*, which are crucial to several adaptive traits in birds. The indel mutations on these genes are not disruptive, and our findings suggest that indel mutations have played an important role in great tit adaptation.

Further investigation is necessary to determine how indels alter protein structure and influence protein function. Structural comparison of MYO5A between the hooded crow and the great tit reveals that an 81-bp great tit–specific deletion extends from one looping region to the beginning of an alpha helix; however, this alteration does not disrupt the structure of the helix itself (Figure 5C). Although structural changes are observed in adjacent regions, the functional and phenotypic consequences of this deletion remain uncertain.

## Conclusion

Indels are common genetic variations, yet our understanding of them remains limited. In this study, we identified indels within coding regions of the great tit genome and analyzed their flanking regions using selection signatures (dN, dS, and *ω*) to investigate the selective forces acting upon them. We show that indels are not only prevalent in coding regions but also subject to positive selection. Moreover, different indel types and lengths are associated with distinct selective pressures, reflecting their heterogeneous characteristics. Additionally, many indels are located in genes that play pivotal roles in adaptive traits, suggesting that indels contribute to adaptive evolution beyond the effects of SNPs in the great tit. Our results highlight that examining the functional and evolutionary impacts of indels provides valuable insights into the genetic basis of adaptation and the mechanisms driving evolutionary change. In line with broader work showing that mutational landscapes, such as CpG distributions, shape coding sequence evolution [43], our findings suggest that indels are an additional and underappreciated component of this landscape.

## Author Contributions

Y.-C.C.: Conceptualization, Methodology, Software, Formal analysis, Visualization, Writing—original draft. J.J.S.W.: Conceptualization, Methodology, Writing—review & editing. T.I.G.: Supervision, Funding acquisition, Project administration, Writing—review & editing.

## Ethical Approval

No ethical issues were involved in this study.

## Data availability

The data supporting the findings of this study are publicly available through the NCBI RefSeq database for the great tit (RefSeq ID: GCF_001522545.3), the collared flycatcher (RefSeq ID: GCF_ 000247815.1), the zebra finch (RefSeq ID: GCF_003957565.2), and the hooded crow (*Corvus cornix*) (RefSeq ID: GCF_000738735.6). The scripts used for data analysis are available on GitHub at https://github.com/chnyuch/pma_indel.git.

## Acknowledgements

This work was supported by the de.NBI Cloud within the German Network for Bioinformatics Infrastructure (de.NBI) and ELIXIR-DE (Forschungszentrum Jülich and W-de.NBI-001, W-de.NBI-004, W-de.NBI-008, W-de.NBI-010, W-de.NBI-013, W-de.NBI-014, W-de.NBI-016, W-de.NBI-022).

## Supplementary Material

Tab.1 *indel_paml* : the selective signatures of indels, insertions, and deletions calculated by PAML with free-ratio and two-ratio models.

Tab.2 *indel_size_paml*: the selective signatures of indels, small indels, and large indels calculated by PAML with free-ratio and two-ratio models.

Tab.3 *indel_list* : indels identified with our pipeline.

Tab.4 *ctrl_300bp*: controls calculated from 300 bp non-indel genes.

Tab.5 *ctrl_600bp*: controls calculated from 600 bp non-indel genes.

Tab.6 *indel_genes*: the gene name and gene ID of indel-harbouring genes.

Tab.7 *indel_enrich_clus*: the clusters from enrichment result of all indels.

Tab.8 *indel_enrich*: the result from enrichment analysis of all indels.

Tab.9 *ins_genes*: the gene name and gene ID of insertion-harbouring genes.

Tab.10 *ins_enrich*: the result from enrichment analysis of insertions.

Tab.11 *del_genes*: the gene name and gene ID of deletion-harbouring genes.

Tab.12 *del_enrich_clus*: the clusters from enrichment result of deletions.

Tab.13 *del_enrich*: the result from enrichment analysis of deletions.

